# Versatile Multiple Object Tracking in Sparse 2D/3D Videos Via Diffeomorphic Image Registration

**DOI:** 10.1101/2022.07.18.500485

**Authors:** James Yu, Amin Nejatbakhsh, Mahdi Torkashvand, Sahana Gangadharan, Maedeh Seyedolmohadesin, Jinmahn Kim, Liam Paninski, Vivek Venkatachalam

## Abstract

Tracking body parts in behaving animals, extracting fluorescence signals from cells embedded in deforming tissue, and analyzing cell migration patterns during development all require tracking objects with partially correlated motion. As dataset sizes increase, manual tracking of objects becomes prohibitively inefficient and slow, necessitating automated and semi-automated computational tools. Unfortunately, existing methods for multiple object tracking (MOT) are either developed for specific datasets and hence do not generalize well to other datasets, or require large amounts of training data that are not readily available. This is further exacerbated when tracking fluorescent sources in moving and deforming tissues, where the lack of unique features and sparsely populated images create a challenging environment, especially for modern deep learning techniques. By leveraging technology recently developed for spatial transformer networks, we propose ZephIR, an image registration framework for semi-supervised MOT in 2D and 3D videos. ZephIR can generalize to a wide range of biological systems by incorporating adjustable parameters that encode spatial (sparsity, texture, rigidity) and temporal priors of a given data class. We demonstrate the accuracy and versatility of our approach in a variety of applications, including tracking the body parts of a behaving mouse and neurons in the brain of a freely moving *C. elegans*. We provide an open-source package along with a web-based graphical user interface that allows users to provide small numbers of annotations to interactively improve tracking results.

## Introduction

Imaging sparse fluorescent signals has become a standard tool for observing neuronal activity. To place that activity in the context of behavior, it becomes increasingly important to perform that imaging in naturally behaving animals (1). Tracking the fluorescent sources through the moving and deforming tissue of these behaving animals is a challenging instance of a multiple object tracking (MOT) problem, and this step is typically a bottleneck for extracting clean measures of activity (2).

Recently, deep learning with convolutional neural networks has been leveraged for many MOT problems with video data including controlling self-driving cars, inferring postural dynamics in humans and animals (DeeperCut (3), DeepLabCut (4), etc. (5)), and computational video editing (non-tracking CGI problems). These advances don’t immediately generalize to videos of fluorescence reported dynamics in living tissue for several reasons.

(1) In contrast to applications like human or vehicle tracking where each object has unique identifiers that can be exploited, two fluorescence signals in the same video are often generated by nearly identical sources and therefore lack distinguishable features (4–7). (2) While transfer learning has been successfully implemented in scientific applications involving natural videos (a horse galloping) (4, 8), the low-level spatial and temporal features detected by these networks rarely reflect structures found in fluorescence microscopy data (9, 10). Thus, this approach rarely reduces the quantity of additional training data required (4, 8, 11–13). Approaches that successfully reduce training data must make hard assumptions about the underlying structure via direct parameter reduction, regularization, or data augmentation (14–17). (3) At the finest spatial scale, convolutional networks rely on images composed of many discriminable textures that typically fill an image (18). Fluorescence microscopy data, however, often has regions of interest with similar fluorescent cells surrounded by voids of black pixels. The combination of sparse global distributions and locally dense homogeneous peaks are less well-suited to convolutional networks, as it becomes harder for convolutional networks to extract useful features for downstream tasks (19, 20). Some methods are proposed to improve the performance of convolutional networks on sparse data but their utility is not shown in the context of MOT (19, 21). (4) Biological videos often exhibit complex motion patterns with nonlinear deformations whereas, in contrast, most vehicle and pedestrian tracking algorithms use linear models or random walks to capture the motion (22).

With sufficiently high frame rates, temporal information can be used to search the vicinity of a cell’s previous location and match identities by minimizing displacement over time. However, motion can often preclude achieving such a frame rate, especially when serially imaging slices of a volume or attempting to recover a signal from a dim fluorescent source. Furthermore, this motion often provides critical context for the problem being investigated (e.g. imaging neuronal dynamics to understand behavior (23)). In these cases, it becomes beneficial to constrain a motion model by maintaining relative positions of cells, correlated motion, and priors for fluorescence dynamics.

Cell tracking methods can be categorized into the following two groups: (1) detect and link, and (2) registration-based. Detect and link algorithms have two distinct steps (6, 11, 13, 14, 16, 24): (1) Detection, where identity-blind candidate locations for objects are proposed by a segmentation or keypoint detection algorithm at each time frame independently. (2) Linking, where temporal associations between detected objects are determined to establish a single continuous worldline across all frames for each individual object. A major drawback of this two-step approach is the propagation of errors from the detection step. Errors that occur in the detection step are difficult to recover from, and they can have detrimental effects on linking and overall tracking quality. Several linking methods have been proposed that are robust to detection outliers, but they either require training with large amounts of manually produced ground-truth data, or are not scalable to lengthy videos (11, 24).

An alternative approach is to directly operate in the image space and optimize some transformation parameters that align a frame to some other frame (4, 12, 17, 25–28). This is done by mapping the underlying image grid from the source to the reference space using the transformation parameters and interpolated pixel values. The transformation parameters must be optimized for each new image over a number of iterations.

Fortunately, recent advances in spatial transformers and differentiable grid sampling have dramatically decreased computational burden and increased performance via GPU acceleration (25, 28–31). Similarly, modern optimization packages such as PyTorch allow the construction of dynamic computational graphs that support more complex nonlinear transformation families and novel cost functions with various regularizers.

Here, we build upon these recent advances to develop ZephIR, a semi-supervised multiple object tracking algorithm with a novel cost function that can incorporate a diverse set of spatio-temporal constraints that can change dynamically during optimization. Our proposed method is capable of efficiently and accurately tracking a wide range of 2D or 3D videos. It allows the user to tune a number of easily interpretable parameters controlling the relative strengths of the registration loss and other constraints, and hence generalizes well to a wide range of biological assumptions. To showcase the efficacy and versatility of our method, we demonstrate its performance on a number of biological applications, including cell tracking and posture tracking.

## Methods

ZephIR tracks a fixed set of keypoints within a volume over time by matching keypoints between an annotated *reference* frame and an unlabeled *child* frame. This matching is done by minimizing a loss function ℒ with four contributions:

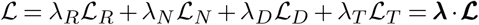

We measure overlap of local image features around the keypoint via ℒ_*R*_. We measure relative elastic motion between keypoints via ℒ_*N*_. We measure the distance of each keypoint to the nearest candidate location from a precomputed set via ℒ_*D*_. We measure smoothness of keypoint-determined dynamical features (e.g. fluorescence or motion) via ℒ_*T*_. Each is described in more detail below.

The relative weights of each term, *λ*, can be freely adjusted by the user to better fit a particular dataset. The user can also set the relative weights to change while tracking a single frame to allow the algorithm to shift focus to different loss components over a number of optimization iterations.

### Image registration, ℒ_*R*_

The first term of our algorithm measures overlap of local image descriptors.

For each keypoint *i* in a child frame, *I*^(*c*)^, an image descriptor (a low-dimensional representation of the local image information), *D*, is sampled according to a sampling grid centered around that keypoint’s coordinates, 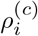. We define a set of parameters, 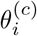, that is closely related to 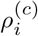 but may include additional transformation models, such as rotation, to characterize the sampling grid, i.e. how each descriptor is sampled from the child frame: 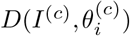.

The descriptors are foveated to prioritize more local information relative to the neighboring features. In lieu of image pyramids (32), we dynamically increase the effective resolution of the descriptors by applying a Gaussian blur at the start of optimization. The blur is decreased in magnitude every few registration iterations. Doing so avoids vanishing or exploding gradients, both of which can occur in regions with sharp, welldefined edges surrounded by a uniform background. On the other hand, restoring the original resolution of the image still provides the best available information for fine-tuning tracking results towards the end of the optimization loop.

Similarly, a set of reference descriptors that serve as registration targets are sampled from a reference frame, *I*^(*r*)^. These are sampled around the user-defined annotations for that reference frame, 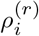, according to a fixed set of parameters, 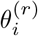.

Using the two sets of image descriptors, our registration loop optimizes the transformation parameters, 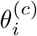, to minimize the following loss term:

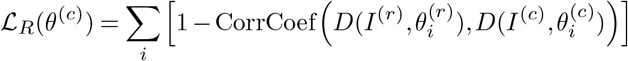

The optimized parameters 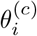 are then used to calculate the desired results, the keypoint coordinates for the child frame, 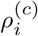. Note that these coordinates are also used for different loss components below, but as 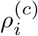 is calculated from 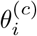, gradients are always accumulated at 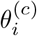.

### Spatial regularization, ℒ_*N*_

Cellular motion within a tissue tends to be highly correlated, but these correlations can be hidden in sparse fluorescent movies that only highlight a small number of cells (or subcellular features) (33). Even in less sparse movies, correlations between nearby keypoints may not be well-captured by descriptors, especially when deformations, noise, or lighting conditions prevent descriptor alignment. In order to reintroduce a similar spatial structure to the data without relying on highly specialized skeletal models, we add an elastic spring network between neighboring keypoints (33–35). The resulting penalty to relative displacement of neighboring keypoints prevents unreasonable deformations, providing a simple and flexible spatial heuristic of the global structure and motion present in the data.

Each of the *i* keypoints being tracked is connected to *j* nearest neighbors to define the following loss term:

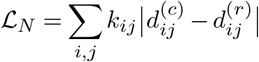

where

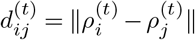

describes the distance between keypoints *i* and *j* in the frame *t*.

When multiple reference frames are available, the stiffness of each spring connection, *k*_*ij*_, is further adjusted to better model the spatial patterns in the data:

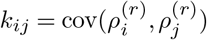

This ensures that connections between highly covariant keypoints are made stronger while connections between keypoints with more weakly correlated motion are weakened or cut accordingly.

### Feature detection, ℒ_*D*_

For this component of the algorithm, we solve an easier problem of identity-blind feature detection, as such detection algorithms have been shown to be fruitful in the context of tracking (6). Namely, we identify key features (such as the center of a cell) present in a volume *without* matching them to a specific feature in some other volume.

This object or feature detection problem has been well-studied, and a wide variety solutions have been proposed. Solutions can range from more parameter-free algorithms (e.g. Richardson-Lucy deconvolution (36, 37)), to algorithms requiring more fine-tuning (e.g. watershed (38)). More recently, deep convolutional neural networks have shown to be powerful, effective solutions as well (e.g. StarDist (10)). Importantly, each of these approaches may work better or worse on different classes of images. Generalization to new datasets can be hard to predict, especially for neural networks that are trained on data generated from a single source.

Our approach is to automatically evaluate simple combinations of these established algorithms by using a shallow model-selecting network. After identifying a set of candidate models, we provide the outputs of these models as input channels to a shallow and narrow convolutional neural network (CNN). If a particular model is best suited for a dataset, network weights for the corresponding input channel are increased during training while suppressing other channels. The low number of learnable parameters in the network also allows fast training for each new type of data or imaging condition, which in turn allows rapid experimentation with new selections of models to test as inputs.

The ultimate output of this selector network, *C*(*I*^(*c*)^), is formulated as a probability map, where each pixel of the original image is assigned some probability of being a desired feature. We use this information to push tracking results towards detected features:

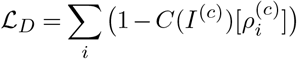

### Temporal smoothing, ℒ_*T*_

Given a sufficiently fast imaging rate, we expect pixel intensity values to be smooth across a small local patch of frames, even for cellular datasets where pixel intensities represent smoothly-varying dynamical signals (39, 40). Thus, we attempt to maintain smoothly-varying local pixel intensities as a form of temporal regularization. For datasets where expected dynamics are appreciably slower than the imaging rate, the strongest version of this regularization is to penalize any deviation from a local zeroth-order fit. We apply this across a small patch of frames (*c* − *ϵ*, …, *c* + *ϵ*) that are registered at once, and add this to the loss for the center frame, *c*:

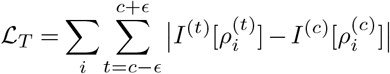

Note that since the loss term is applied for the center frame only, it does not affect the results for the other frames despite registering all frames in the patch together. Additionally, this component of the algorithm requires registration (or approximate registration) of nearby frames, making it more appropriate in low-motion conditions or after initial coarse registration is complete.

### Frame sorting

Using all or some of the loss terms listed above, a single *child frame* is registered to a *reference frame*, and all keypoints in the frame are tracked simultaneously. To fully analyze a movie, we need to register every frame to a reference frame.

For many datasets, it is best to register every child frame directly to a coarsely similar reference frame, and let annotations for that reference frame provide initial guesses for keypoints in the child frame (Fig. 2A). For this, we must identify a set of representative reference frames that capture the range of deformation patterns present in the movie, and we must assign each remaining frame to one of those reference frames. A pairwise distance between all pairs of frames is determined by some similarity metric (e.g. correlation coefficient) applied to low-resolution thumbnails. A k-medoids clustering algorithm is applied to these pairwise distances to identify a small number of median frames to best serve as reference frames for all other frames in the corresponding cluster (Fig. 2) (4, 26).

In other datasets, the registration results from one frame in a cluster may provide useful insight into the solution for a different frame in that cluster. For example, a frame that is close (in deformation space) to the reference may be easy to track. The tracked results from that frame, in turn, may provide a better guess for keypoints in a frame that is further away from the reference. This can reduce the distance between the initial guess and correct positions, and thus reduce the difficulty of the optimization problem. Thus, every child frame being registered is associated not only with a reference frame (a registration target), but also a previously registered *parent frame*, which provides the initialization prior to optimization (Fig. 2B).

Additionally, the learning rate for the child frame is partly determined by the distance between the parent and child frames. We expect that when a parent-child pair are close in the deformation space, the keypoints do not undergo significant local displacements. Hence, a low learning rate is applied for a similar parent-child pair, scaling up to a high learning rate in the case of a dissimilar pair to allow tracking of features much further away. The combination of these effects produces a flexible limit on the range of possible optimization results for the child frame based on coarse similarity to its parent frame (41–43).

To take full advantage of this parent-child interaction, we sort all frames into distinct sequences of parent-child frames based on similarity. Each of the resulting *branches* begins from a previously selected median reference frame. The subsequent child frames are selected to minimize the distance from a parent frame until every frame is assigned to a branch. Doing so produces unique sets of frames that stem from each reference frame, naturally forming clusters that separate similar frames from dissimilar ones. This is particularly useful for datasets that repeatedly sample from a limited set of postures or global spatial structures (e.g. locomotion).

However, not all datasets have temporal patterns that can reliably make use of the similarity-based initialization method. For such datasets, a chronologically sorted queue may be more reasonable and provide better accuracy overall, where a branch simply stems from each reference frame both forwards and backwards in time until it encounters the first frame, the last frame, or another branch (Fig 2). Note that the parent-child interactions during tracking are still the same regardless of the sorting method. For a chronologically sorted queue, the controlled variation of learning rates effectively allows us to adapt to different capture frame rates. A high frame rate video often captures smooth motion that benefits from low learning rates but a low frame rate video does not.

#### Algorithm 1

ZephIR optimization loop

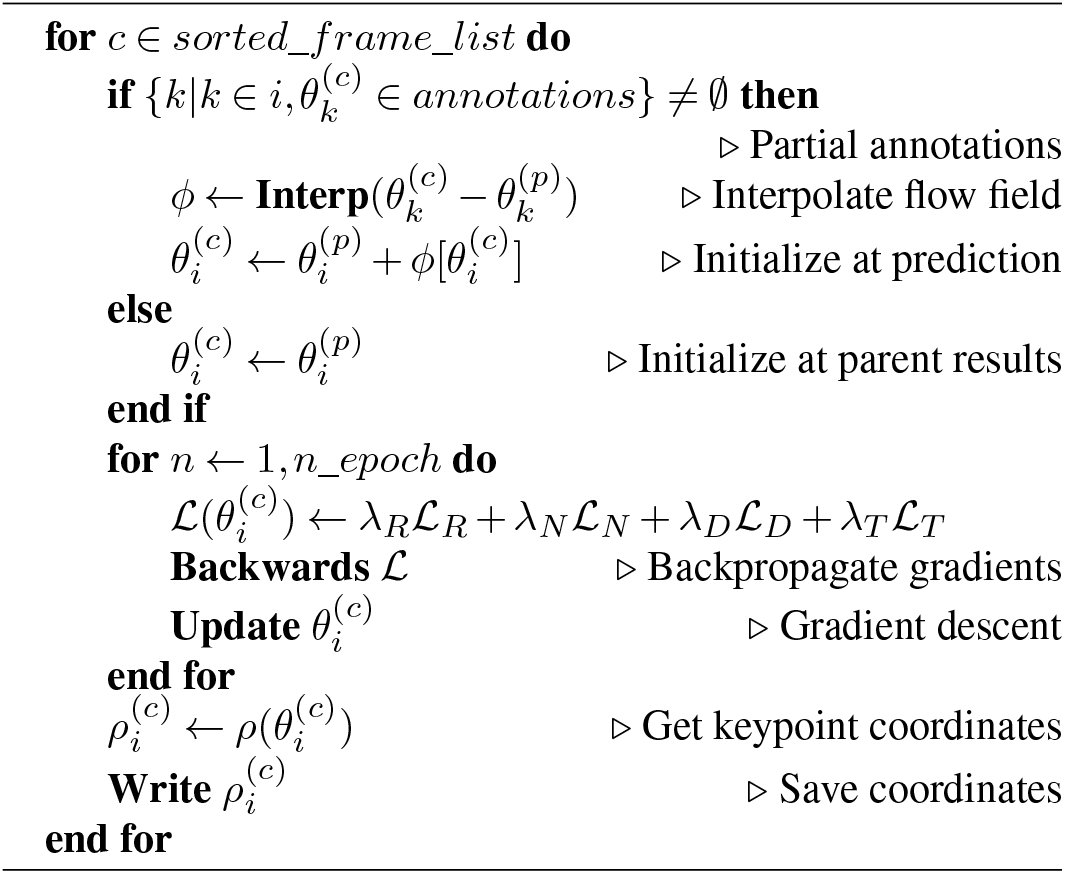

### User intervention

Our pipeline allows a user to dramatically improve tracking quality in various ways by providing further supervision. Providing additional fully annotated frames will improve registration targets to better match descriptors from similar frames. Strategically selecting a new reference frame can have dramatic impacts on frame sorting as well, creating opportunities to form tighter clusters of parent-child branches.

Furthermore, when multiple reference frames are present, covariance of keypoints in those frames helps better define an implicit global spatial structure by modulating stiffnesses of the spring connections between neighboring keypoints, *k*_*ij*_. Any additional reference frames can provide more accurate covariances, and thus a spatial model that is more accurately tailored for that particular dataset.

*Partially* annotated frames are not used to seed sorted frame branches nor used to sample reference descriptors. Still, all user annotations present in the frame are utilized to improve the tracking quality of the remaining keypoints in that frame (Fig. 1D).

**Figure 1.**
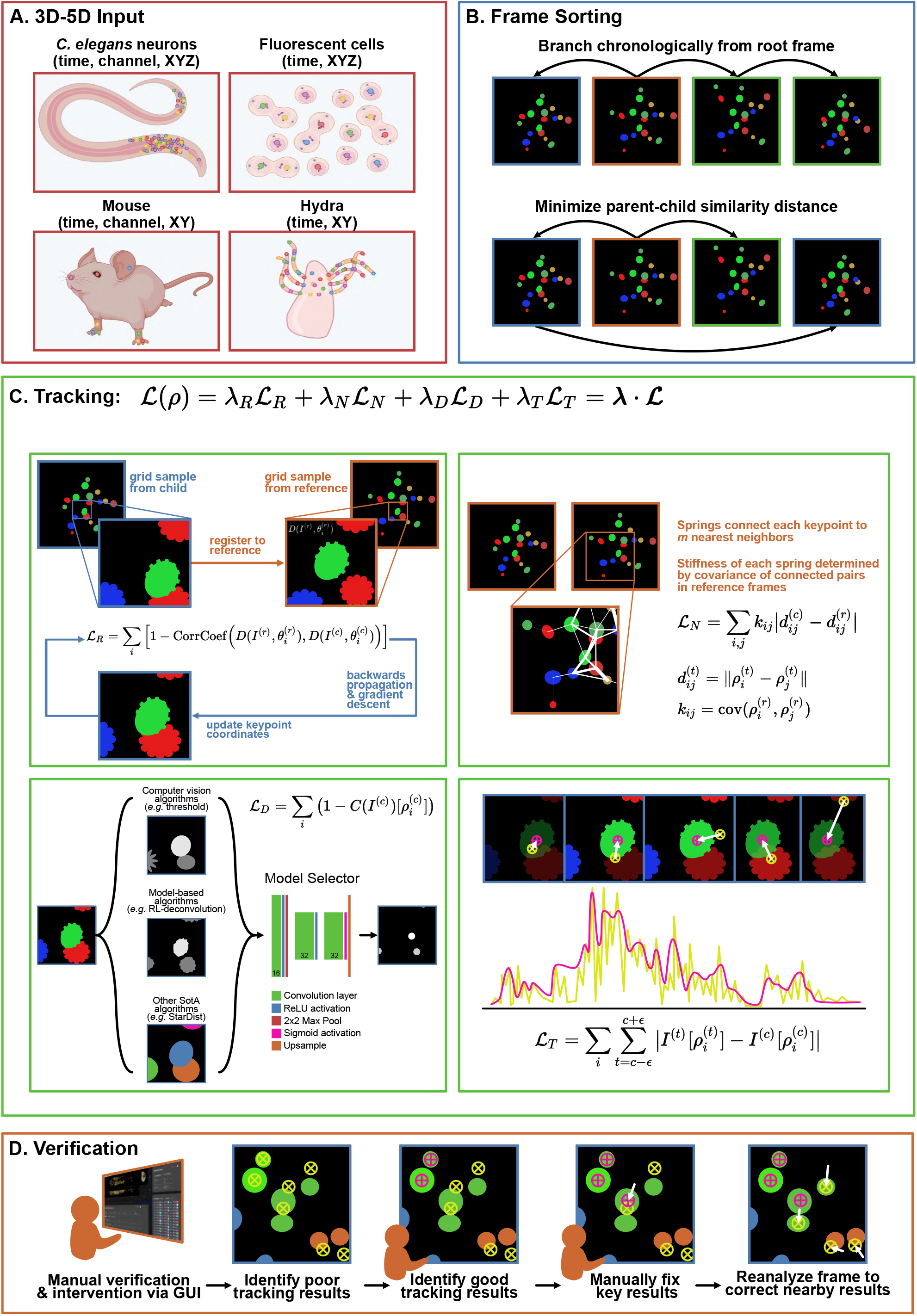
Overview of ZephIR algorithm. **A**. Examples of input datasets. ZephIR can track keypoints in various biological systems, including fluorescent cellular nuclei in a tissue and body parts that summarize a posture. Input dimensions can range from 3D (time, XY) to 5D (time, channel, XYZ). Colored dots indicate example keypoints to be tracked. **B**. Frame sorting schemes. A branch defines an ordered queue of frames to be tracked. Each branch begins at a manually annotated reference frame (orange), **B**. but subsequent parent (blue) and child (green) frames in a single branch can be sorted either by chronology (top) or by minimizing the similarity distance between each parent-child pair (bottom). **C**. Overview of tracking loss. Tracking loss is comprised of four terms: 1) overlap of local image features around each keypoint, sampled from the current frame and its nearest reference frame, 2) elastic connections between neighboring keypoints with varying stiffnesses based on covariance of the connected keypoints, 3) proximity to features detected by a shallow model selector network that takes in a number of existing feature detection software as input channels, 4) smoothness of temporal dynamics at each keypoint position. **D**. Overview of steps for manual verification and additional supervision. Users can verify tracking results as correct or identify incorrect results. After fixing a few key incorrect results, ZephIR can use those new annotations as well as the verified correct tracking results to improve tracking results for all other keypoints in that frame (and all its child frames).

Firstly, prior to gradient descent, displacements between all available annotations and their corresponding coordinates from the parent frame are used to interpolate a flow field. This flow field serves as a rough model of the global motion between the two frames (17, 27, 35). We sample from the flow field at the remaining keypoints coordinates in the parent frame and apply the resulting estimated displacements to initialize the keypoints closer to their new positions in the child frame. This is particularly helpful for pairs of parent-child frames with large motion between them, and the flow field can always be improved in both precision and accuracy by adding more annotations for the child frame.

Secondly, the spatial regularization during the optimization process, ℒ_*N*_, also makes good use of any partial annotations. The annotations are fixed in place, but the spring connections to their neighbors remain a crucial component of the backwards gradient calculations and helps to “pull” the connected keypoints into place.

To streamline the process of providing user supervision, we offer a browser-based graphical user interface that provides an intuitive, simple environment to produce and save further annotations. Since our approach lacks a slow “training” phase, any new annotations can be applied to tracking a frame directly from the GUI. A macro available in the GUI executes a temporary state of the algorithm quickly and efficiently, allowing users to see the precipitated improvements immediately.

Additionally, the GUI provides an opportunity for users to provide supervision without creating new annotations. The user may upgrade individual results into annotations or entire frames into new reference frames by marking them as correctly tracked. These user-confirmed frames will be treated as a regular reference frame next time the algorithm is executed, benefiting from all the improvements to tracking quality discussed previously. These improvements to the rest of the results can be observed immediately by executing the algorithm from the GUI.

## Results

### Neurons in crawling worms (C. elegans)

Optical methods based on fluorescence activity of calcium binding indicators has become a standard tool for observing neuronal activity in *C. elegans*. To do so, it is necessary to track fluorescent signals from individual neurons across every frame in a recording. This poses a significant challenge, particularly when the animal is allowed to freely crawl. The worm’s brain undergoes fast, dramatic, nonaffine deformations, exhibiting a large variety (forward and backward motion, omega turns, coils, pharyngeal pumping, etc.) and magnitude (up to ten microns relative to an internal reference frame) of movements as the animal behaves (22, 23, 44, 45).

Many solutions have been proposed to track fluorescent neurons in *C. elegans*. Two step (detect and link) approaches often suffer from the lack of reliable detection algorithms and require relatively low frame-to-frame motion in order to accurately link the detected neurons (6, 12, 16). Similarly, deep learning approaches are limited by insufficient training data, often failing to generalize across different animals, even those within the same strain (11, 26, 46). While these approaches have provided important insight and progress, there remains substantial need for improvement in accuracy and efficiency when tracking many neurons in freely behaving worms.

Fig. 3 describes the workflow and performance of ZephIR on tracking a set of 178 neurons in the head of a freely behaving worm across a recording of approximately 4.4 minutes (1060 frames @ 4Hz). The video has been centered and rotated to always face the same direction, but no further straightening has been done. With only a few manually annotated reference frames, ZephIR already achieves state-of-the-art MOT accuracy (20, 47, 48) as reported on similar datasets in recently published works (11, 12, 16) (Fig. 3A,B).

**Figure 2.**
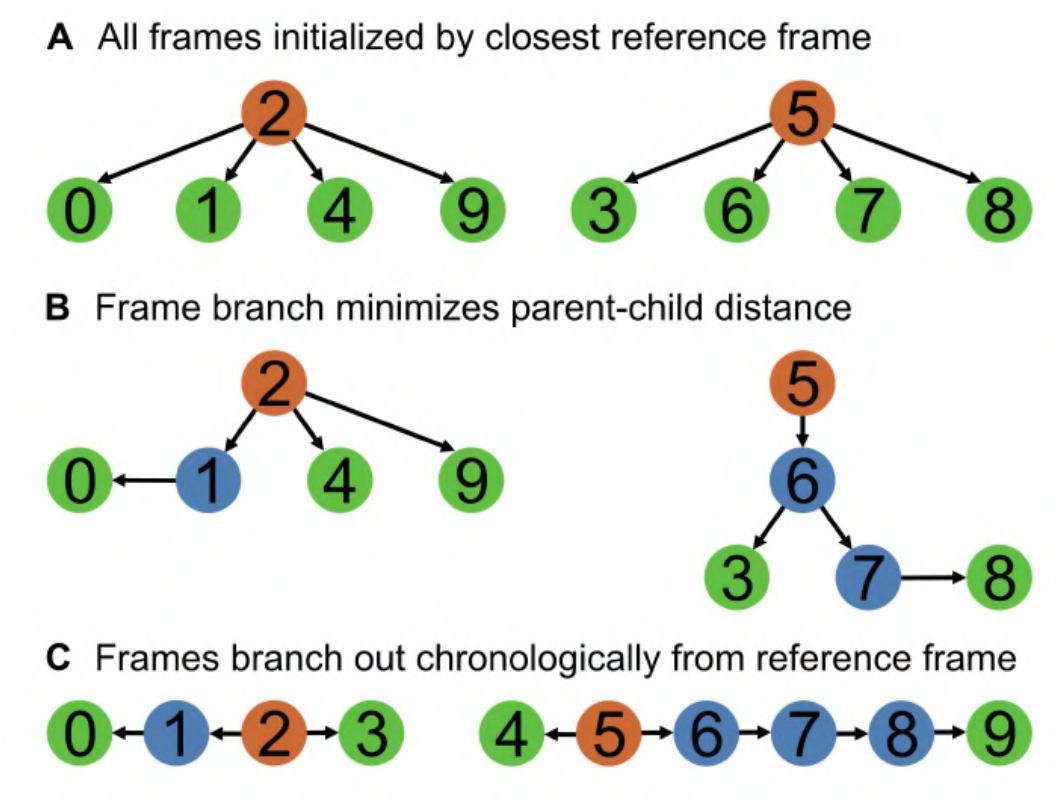
Overview of frame sorting strategies. Orange indicates fully annotated reference frames, blue indicates parent frames with at least one child frame, and green indicates child frames. **A**. In the simplest strategy, all frames are initialized by the closest reference frame. **B**. Frames are sorted into ordered queues based on similarity. Each of these branches start with a reference frame, and new child frames are added such that the parent-child similarity distance is minimized, naturally clustering similar frames around each reference frame. **C**. Frames are sorted chronologically, branching both forward and backwards from each reference frame.

**Figure 3.**
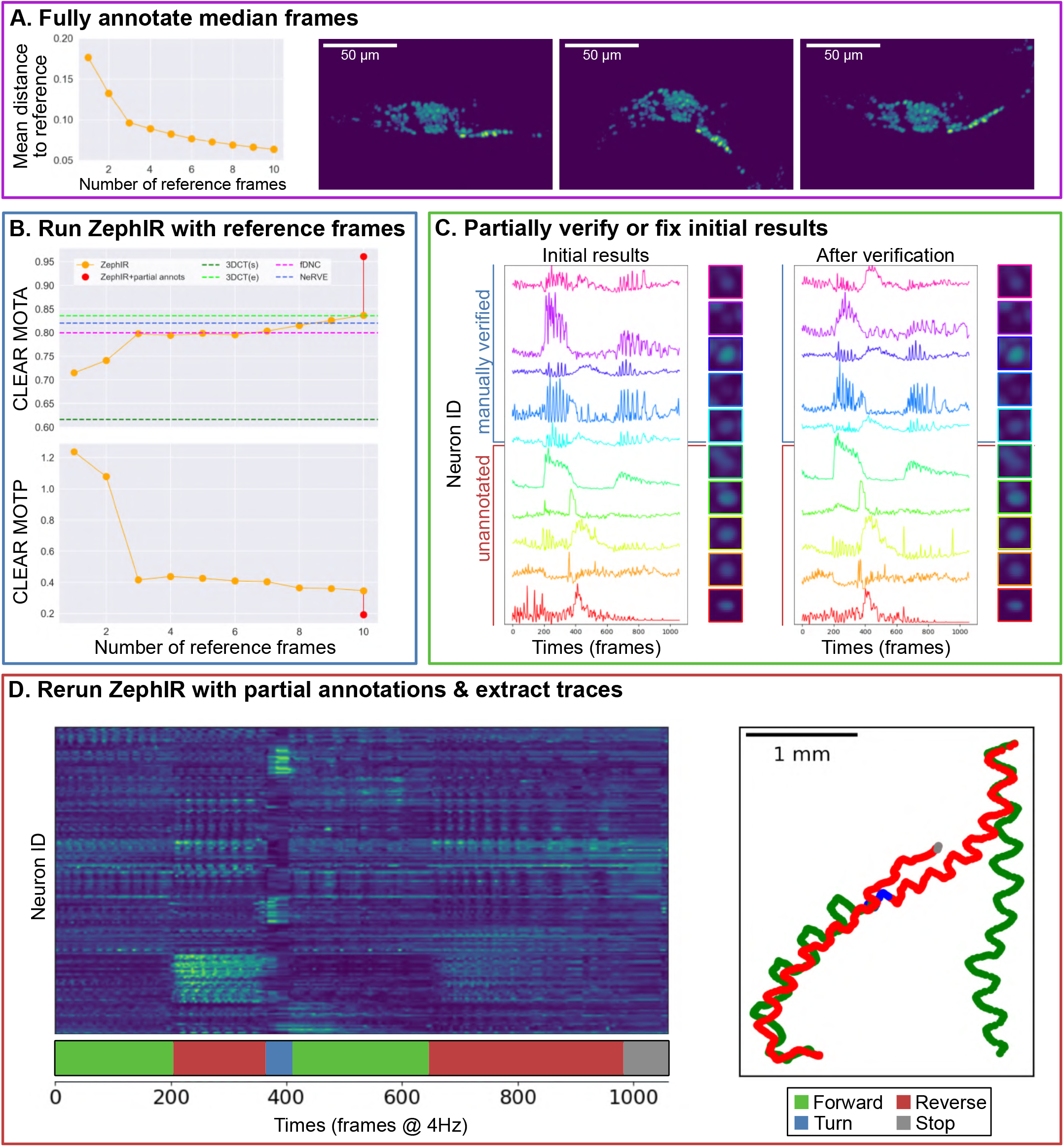
Results for freely behaving *C. elegans*. **A**. Plot of mean distance to the nearest reference frame vs the number of reference frames (left), and the first three median frames recommended by ZephIR’s k-medoids clustering algorithm (right). The first three median frames clearly represent the three main postures that the worm cycles through as it crawls. **B**. MOT accuracy (higher is better) and precision (lower is better) vs the number of reference frames. Note that once the majority of the postures present in the data is well-represented by the first three reference frames, subsequent additions returns diminished improvements. Last data point shows ZephIR’s accuracy using 10 reference frames with 10 partial annotations across all frames (panel C). We also compare ZephIR’s accuracy with Neuron Registration Vector Encoding (NeRVE) (16), fast Deep Neural Correspondence (fDNC) (11), and 3DeeCellTracker (12) in both single (3DCT(s)) and ensemble (3DCT(e)) modes as reported in their respective publications. Note that the accuracies from 3DeeCellTracker reflects both errors in detection and tracking. **C**. 10 neurons were randomly selected to be verified or corrected to serve as partial annotations. Traces of 5 of these neurons extracted using the initial ZephIR results with 10 reference frames (left), and those using verified true positions (right) are shown, along with 5 other randomly selected neurons. Traces are calculated as fold change over the baseline, where the baseline is defined as the intensity in the first frame. Tracking quality for these 10 neurons can also be seen in individual crops around the neurons averaged across all frames (sharper image of the cell at the center reflects better accuracy and precision in tracking). Note how the five unannotated neurons show improvements in tracking quality after the addition of partial annotations, exemplifying the effects of partial annotations on the unannotated neurons in the same frame. **D**. Neuronal activity traces from 178 neurons, extracted using results from ZephIR with 10 reference frames and 10 partial annotations in all frames. Traces are calculated as fold change over the baseline, where the baseline is defined as the intensity in the first frame. Behavior is shown in the ethogram below the heatmap. Trajectory of the worm (t=0 at bottom right) is also colored with the behavior state at the time. Trajectory of the worm matches changes in behavior over time as expected, and many of the neuronal activity traces show strong correlation with behavior.

We further improve on the accuracy of the initial results by providing additional supervision. We randomly selected ten neurons uniformly distributed throughout the brain to verify and use as partial annotations across all frames. Because the initial results already achieved high accuracy, they only required correction for a subset of frames (≈ 15%). After this correction and validation, annotations for these ten neurons were re-classified as manual annotations in all frames. The partial annotations produce a dramatic improvement in accuracy (red data point in Fig. 3B) without the need to verify entire frames.

Through this workflow, we are able to achieve a sufficiently high accuracy to extract good, meaningful neuronal activity traces across the entire recording (Fig. 3D) (39, 40). Many neurons show clear correlation with observed behaviors, and the activity patterns are comparable to previously published works (16, 22, 49, 50).

### Detect-and-link tracking for multimodal images

In multimodal images, directly comparing descriptors from different modalities typically will not generate useful gradients for image registration. Nonetheless, other terms in our loss can be used as the “link” step of a detect-and-link algorithm to associate detected keypoints between modalities (ignoring all image information). As an example, we linked *C. elegans* neurons between a bright-field image of a worm and a slightly deformed fluorescent image of nuclei in the same animal (as in the previous section) (Fig. 4A). We provided ZephIR with nuclear positions in each image, and show that it is able to utilize the other loss terms to link keypoints between the two (Fig. 4).

**Figure 4.**
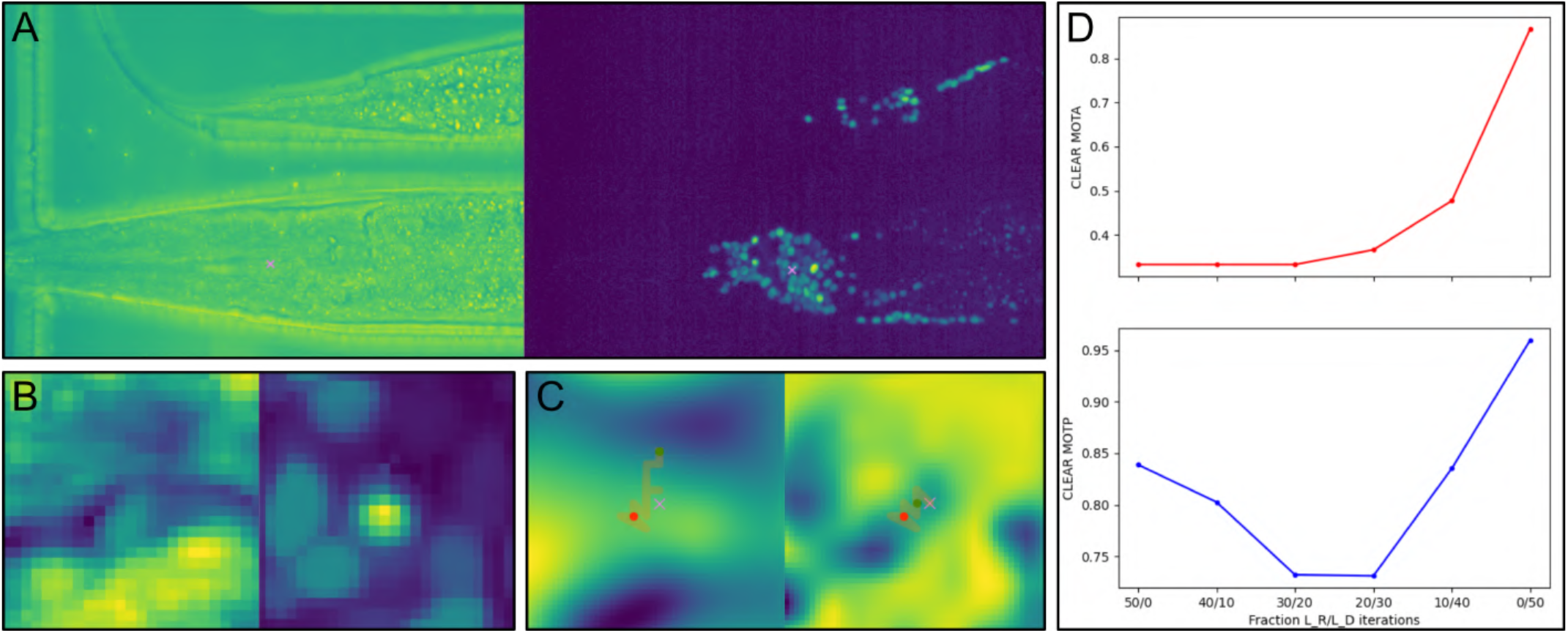
Results for multimodal images of *C. elegans*. **A**. Two frames in different imaging modes. The reference frame is a widefield image of the worm body (left). The child frame is an image of GCamP fluorescent nuclei of neurons in the worm’s brain (right). **B**. Image descriptor used to register (ℒ_*R*_) an example neuron as sampled from the reference volume in widefield mode (left) and from the child volume in GCamP mode (right). **C**. Visualization of the registration loss (ℒ_*R*_) between the two descriptors in panel B (left), and of the feature detection loss (ℒ_*D*_) for the child frame (right). The optimization trajectory (orange) is displayed on top, starting from the initial coordinates (red) and ending at the optimized coordinates (green). Purple cross indicates the ground-truth coordinates. When using registration loss only (left), optimization fails to find the correct coordinates due to a lack of any local minima nearby. In contrast, when using feature detection loss only (right), optimization is able to find the basin around the correct coordinates. **D**. ZephIR MOT accuracy and precision for the child frame. A total of 50 optimization iterations are separated into two phases: a registration phase with *λ*_*R*_ at 1.0 and *λ*_*D*_ at 0.0, and a feature matching phase with *λ*_*R*_ at 0.0 and *λ*_*D*_ at 1.0. To tune the relative contributions of the two losses for tracking, we vary how many iterations are given to each phase. Contributions from other loss terms, *λ*_*N*_ and *λ*_*T*_, are kept constant for both phases. Despite the low accuracy when using registration phase only (left), the feature matching phase is able to recover tracking accuracy (right).

Note that ZephIR requires a separate algorithm (or manual input) to detect keypoints in each modality to do so. This detection can be performed with the built-in model selector approach (*L*_*D*_ in Fig. 1C) or any user-provided algorithm.

### Posture of a behaving mouse

Here, we demonstrate how ZephIR can be used for behavioral tracking in natural movies by analyzing the pose of a head-fixed mouse performing a motor task. The richness of local image features present in natural images lend themselves to registration. In addition, by connecting key points along the mouse’s body, our spring network loss (ℒ_*N*_) can implicitly capture the scaffold underlying the mouse’s posture.

There exist many solutions for similar problems in posture tracking. In particular, convolutional neural networks have been successfully implemented for posture analysis in both laboratory and natural settings (3–5). Notably, DeepLabCut adapts a ResNet CNN architecture to track postures of various animals without any physical markers. DeepLabCut utilizes transfer learning, where a “base” model is trained on a publicly available dataset of various “natural” images prior to specializing the weights to a particular dataset. Their method reduces the amount of training data required to achieve state-of-the-art results by orders of magnitude (from hundreds of thousands to just a couple hundred labeled images), and thus reduces the amount of manual labor required by the experimentalist.

Fig. 5 compares the performance of our algorithm and that of DeepLabCut on the same dataset. We track ten points that summarize the mouse’s posture as it performs a task. We show that for low numbers of reference frames, i.e. low numbers of training data, ZephIR can produce much better quality tracking than DeepLabCut, achieving good results with less than 20 reference frames. ZephIR is also able to produce this result with much less total computation time as it does not require a slow training phase.

**Figure 5.**
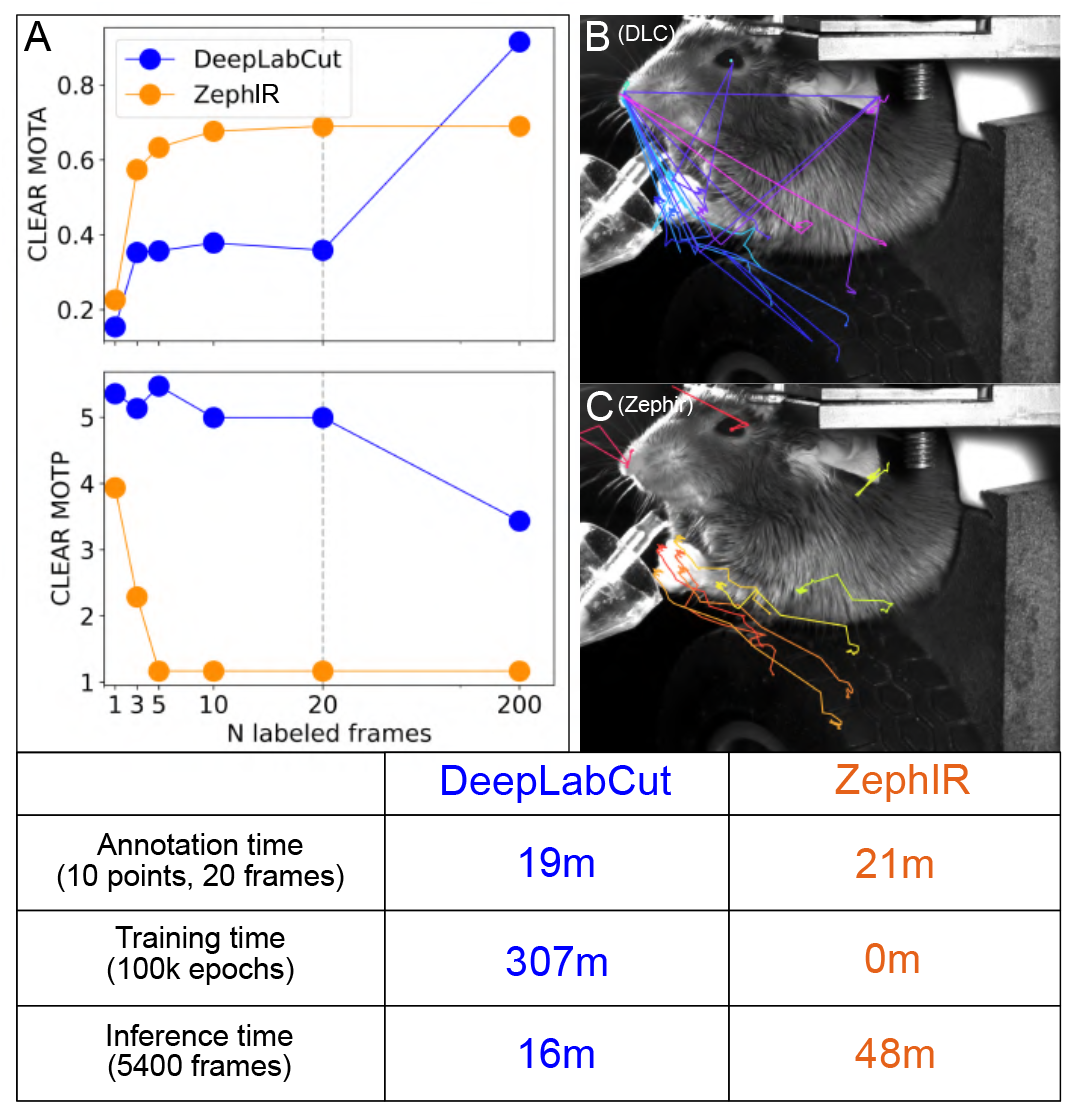
Results for a behaving mouse. We compare performances of ZephIR and DeepLabCut on tracking 10 body parts that characterize the mouse’s posture over time. **A**. MOT accuracy and precision vs the number of manually labeled or ground truth frames. These labeled frames are used as reference frames for ZephIR and as training data for DeepLabCut. The frames are selected based on automated recommendations from each algorithm, meaning the two sets of frames used may not be identical. The last data point (200 labeled frames) for DeepLabCut are produced with training data generated by verifying and correcting ZephIR results with 10 reference frames. Note that ZephIR achieves better accuracy when only a few labeled frames are provided, but DeepLabCut ultimately reaches a higher accuracy when its training data was augmented with ZephIR. **B, C**. DeepLabCut and ZephIR results with 20 labeled frames (vertical line in panel A) for tracking mouse body parts as it raises its paws. Note that ZephIR is more stable during motion while DeepLabCut tends to jump between the different body parts. **Table**. Annotation and computation speed comparison. Annotation time is calculated for the same person, using the respective GUI’s provided with each software package. Training and inference times are tested on the same CPU and single GPU environment and with 20 reference frames (vertical line in panel A). While DeepLabCut is faster for inference, it requires a slow training phase, dramatically increasing the total computation time.

It is important to note that DeepLabCut can ultimately produce more accurate results when provided with more training data. However, in the case that an experimentalist requires higher accuracy than ZephIR is able to provide on its own, ZephIR can easily fit into a DeepLabCut workflow to augment the amount of training data available. Instead of manually labeling the full list of 200 frames to produce the last data point in Fig. 5A, we only annotated the first 10 of the recommended frames. We then run ZephIR using those frames as references, verify the tracking results for the remaining 190 frames, and correct any errors to produce the full set of 200 training images to use for DeepLab-Cut. Including this step dramatically cuts the total human time required for a DeepLabCut workflow, from an extrapolated 160 minutes to label all 200 frames by hand to 53 minutes.

More recently developed variants of DeepLabCut, such as Deep-GraphPose, can also reduce training data size by incorporating spatio-temporal priors and enabling semi-supervised training that uses both annotated and unannotated data (5). However, these variants still require a significant amount of training data (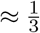 of DeepLabCut’s requirements, compared to ZephIR’s ≈ 5 − 10%) and often fail when analyzing sparse or volumetric datasets, making it difficult to employ for biological datasets.

### Performance

We are able to compute the loss terms and optimize the tracking parameters efficiently by utilizing modern deep learning tools with, in particular, differentiable grid sampling and GPU acceleration offered by PyTorch (51). Since our approach does not require a training phase, which is often the most significant bottleneck in both time and resources, it is fast without being computationally costly. We also sacrifice a small amount of performance to reduce the amount of memory required for both CPU and GPU to levels that are reasonable for commercial laptops. This balance can be manually adjusted by the user depending on their computing environment.

We run the following tests on a PC with a 16-core AMD Ryzen Threadripper 1950X processor @ 3.40GHz, 64GB RAM, and an Nvidia GTX 1080Ti GPU, 11GB VRAM. The tests were carried out on the freely behaving worm dataset (Fig. 3).

In the default configuration, ZephIR registers 100 1×5×25×25 (CxDxHxW) descriptors (ℒ_*R*_) with spatial regularization (ℒ_*N*_) over 40 optimization epochs for an average of 1.24s total computation time spent per volume. In comparison, similar algorithms such as NeRVE takes an approximate 50sec/vol on over 200 computing cores (16) and 3DeeCellTracker approximately 2min/vol on a desktop PC with an NVIDIA GeForce GTX 1080 GPU (inference only) (12).

During the test, the process utilizes a maximum of 1.84GB RAM and 0.89GB VRAM. The number of descriptors does not significantly affect performance as descriptors are registered in parallel, but the number of epochs will impact speed linearly. The size of descriptors may slightly affect performance as well as memory consumption.

## Discussion

ZephIR is a semi-supervised multiple object tracking algorithm. It tracks a fixed number of user-defined keypoints by minimizing a novel cost function that dynamically combines image registration, feature detection, and spatio-temporal constraints. Local registration of image features enables tracking of keypoints even in sparse imaging conditions, such as fluorescent cellular data, while a spring network incorporates a flexible motion model of the neighboring keypoints without the need for a highly specialized skeletal model. Feature detection can help fine-tune tracking results to match a nearby detected feature in the image or even recover good tracking accuracy in cases where registration clearly fails to produce good gradients. The model utilizes modern deep learning libraries, recent innovations in spatial transformers, and optimization tools to calculate loss and backpropagate gradients efficiently in a GPU environment.

We demonstrate that our approach is able to reach state-of-the-art accuracy on a diverse set of applications, including extracting neuronal activity traces in a freely moving *C. elegans* and tracking body parts of a behaving mouse. Notably, ZephIR is able to do so with a small amount of ground-truth data and low computational resource requirements. Recent deep learning-based methods often require large amounts of labeled frames for each new dataset. In contrast, ZephIR is able to generalize to radically different datasets with just a few labeled frames and adjustments to some hyperparameters.

Any amount of new manual labor, whether simply verifying correct results or fixing incorrect ones, can dramatically improve ZephIR’s accuracy. Verifying or correcting entire frames produces new reference frames to provide better reference descriptors for registration and improve flexibility of the spring network. Verifying only a subset of keypoints can initialize better tracking guesses for all other points in the same frame by interpolating a global motion model between parent and child frames. Additionally, any improvements in tracking a frame can cascade down to all its child frames, further reducing the amount of supervision required.

Through this workflow, ZephIR achieves unprecedented accuracy with minimal manual labor, even on a freely behaving *C. elegans*, where large deformations present a challenging tracking problem. We also expect to achieve similarly strong performance on sparse fluorescent videos of deforming neurons in other models organisms including *Hydra*, zebrafish, and *Drosophil* (1, 46, 52, 53).

With its versatile design and low computational requirements, ZephIR is designed to be highly accessible and useful for a diverse set of applications. On the other hand, we hope to also support full utilization of more powerful computational environments, especially when multiple GPUs are available. In particular, since distinct frame branches do not interact with one another when tracking, we may split them across multiple machines or GPUs to analyze in parallel, resulting in roughly linear gains in speed. These performance gains could be available to all users by hosting an updated version of our annotator GUI on a dedicated GPU server.

A notable limitation of our approach is that at least one annotated frame is required. We hope to mitigate this issue through future key upgrades. For example, we hope to use an object detection algorithm to automatically annotate the first reference frame, where linking or identity-classification is not necessary (7, 10, 15, 31, 54). Many experiments with immobilized animals or low-motion data often only need one reference frame, meaning such datasets could be tracked entirely unsupervised. Advancements in spatial transformers and novel motion models may also eliminate or reduce the need for partial annotations to initialize keypoint coordinates closer to their true positions than the parent coordinates alone (2, 17, 25).

For some datasets, other approaches may be more accurate than ZephIR. As the field of deep learning continues to develop, we can expect more powerful, generalizable models to emerge. Still, ZephIR can be a powerful data augmentation tool upstream of any of these algorithms, as was demonstrated with behavioral mouse data in this work. Since it can reach reasonable accuracy with a low number of annotations, ZephIR can reduce the amount of labor required to produce the necessary training data. It may be a key component in generating a critical amount of ground-truth data to build new models to perform multi-object tracking in particularly challenging datasets.

ZephIR is available at:

https://github.com/venkatachalamlab/ZephIR.

## Supplement

### Loss visualized

#### Verifying frame tree construction

ZephIR builds a tree with each branch forming an ordered queue of frames to be tracked. Each tree begins at a reference frame and follows a sequence of parent and child frames. Methods for selecting “optimal” reference frames for a dataset and new child frames for a branch were determined heuristically and then verified. Fig. 7 tests our reference frame selection method and the tracking accuracy for all other frames produced with those reference frames. Fig. 8 tests our child frame selection method and the tracking accuracy for candidate child frames given a single parent frame. In both cases, we can verify a strong correlation between the score used in our selection method and the tracking accuracy.

**Figure 6.**
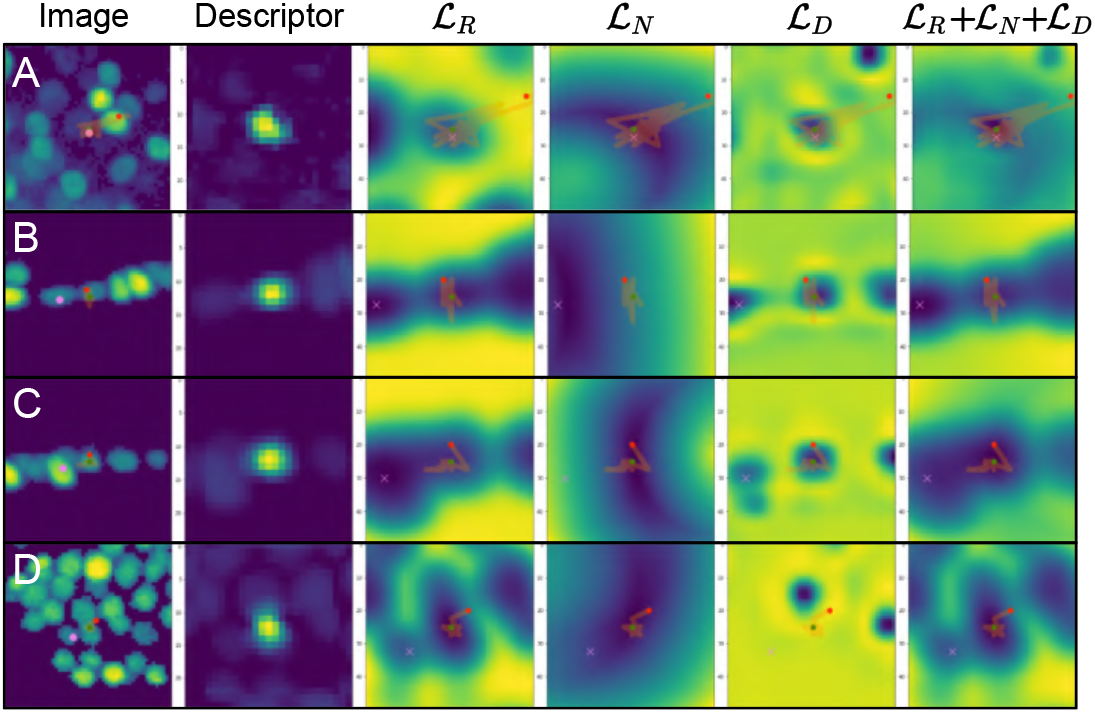
Visualization of the loss map around certain neurons in freely moving *C. elegans* with all other neurons fixed at the ground truth position. Columns, from left to right: visualization of the volume around the neuron, the image descriptor of the neuron used for registration, map of the registration loss, map of the spring network loss, map of the feature detection loss, and map of the sum of the three previous losses. First column is centered at the initial coordinates, all others at the final optimized coordinates. Each row analyzes a different neuron and its optimization trajectory: initialized at red, optimized along orange, final results at green. Purple marks the ground truth position for the neuron. Comparing different loss maps along with the overall optimization trajectory can help diagnose certain tracking issues and give valuable insight on how to optimize the loss weights, *λ*, for a particular problem. **A**. For this neuron, all three loss components provide good minima at the ground truth position. ZephIR easily finds the correct result through gradient descent. **B**. For this neuron, ZephIR fails to escape a local minima present in both registration loss (ℒ_*R*_) and feature detection loss (ℒ_*D*_) at a neighboring neuron. However, we can see that spring network loss (ℒ_*N*_) creates good gradients that could push the neuron out of initial basin, thus increasing *λ*_*N*_ may improve this result. **C**. For this neuron, ZephIR fails to escape a local minima at a neighboring neuron. In contrast to row B, the spring network loss (ℒ_*N*_) contributes to this local minimum, but the registration loss (ℒ_*R*_) provides gradients towards a basin at the correct position. Thus, decreasing *λ*_*N*_ may improve this result. **D**.. For this neuron, all three loss components fails to present global minima at the ground truth position, and only the registration loss (ℒ_*R*_) presents a local minimum there. Since the neuron position is initialized such that it must cross a deeper minimum to reach the correct position in all loss maps, adjusting *λ* alone may not be able to improve this result.

**Figure 7.**
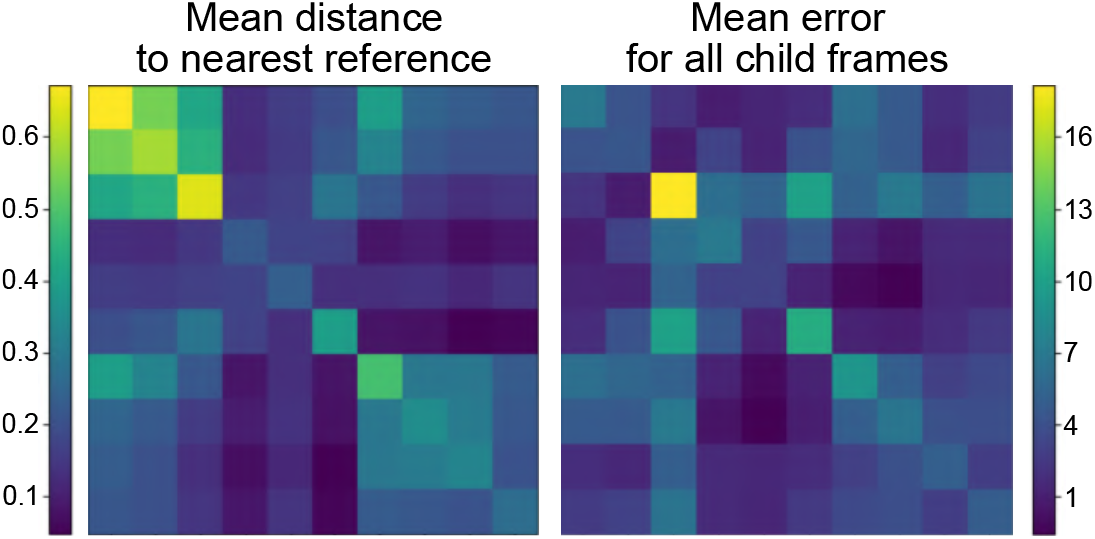
Verifying reference frame recommendation. ZephIR recommends frames to annotate as reference frames via k-medoids clustering of low-resolution thumbnails. We test ten candidate frames and all pairwise combinations. During clustering, each child frame is assigned to a cluster around a reference frame based on minimum distance (i.e., assigned to nearest reference frame). With each update, the clustering minimizes a score based on the mean distance between child frames and their assigned reference frame (left, lower/bluer is better). The results from tracking with the two candidate reference frames are evaluated for all other frames (right, lower/bluer is better). We can compare the resulting profiles of score and accuracy for each pair of candidate frames in order to evaluate the efficacy of the recommendation method (more similar is better).

**Figure 8.**
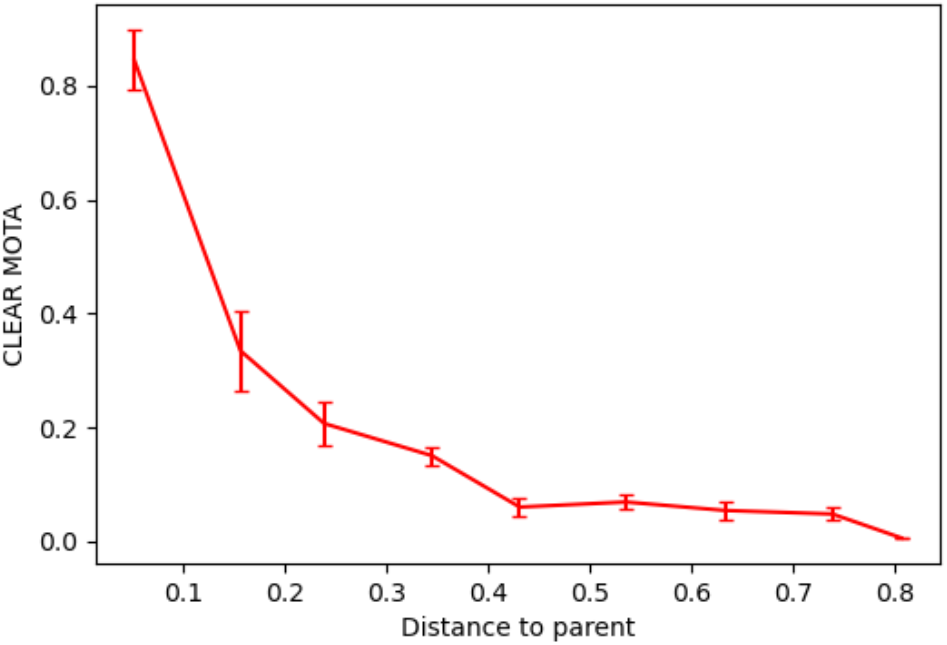
Verifying parent-child selection. When sorting based on frame similarity, each subsequent child frame is selected to minimize the distance from a parent frame. We test pairs of frames to study the effect of distance between parent and child frames. In these tests, the parent frames provide both the initial positions of the keypoints in the child frame and the reference descriptors as registration targets, and tracking results for keypoints in the child frame are evaluated. It is evident in the resulting curve that accuracy quickly falls with distance.

In addition to the current implementation, we explored several other avenues in which parent and reference frames may improve tracking quality for the child frame.

Notably, target descriptors for image registration are currently sampled from a single reference frame at the root of the frame branch. We tested methods that made direct modifications to image descriptors based on parent results, including:

1. sampling target descriptors from the parent frame
2. deforming target descriptors sampled from reference frames based on flow fields between parent and reference frames
3. deforming child descriptors based on flow fields between parent and child frames
4. averaging samples from all preceding frames in the branch to create target descriptors.

We also tested other implementations of motion prediction (for better initialization of keypoint coordinates for the child frame), including models based on:

1. momentum of keypoints across preceding frames in the branch
2. piecewise global deformation fields between parent and child frames fit prior to tracking keypoints
3. two-step tracking, where results from the first iteration of tracking would be used to generate a low-frequency global deformation field between parent and child frames

Unfortunately, all of the above created a tighter relationship between parent-child pairs that was ultimately too sensitive to errors and error propagation down the branch.

We also explored potential ways to use all available reference frames for tracking a single child frame. In particular, we tested simultaneous registration to target descriptors sampled from each of the available reference frames. We implemented this system in two different flavors:

1. target descriptors from different reference frames are stacked along an axis as separate data channels, and registered with copies of the child descriptors, producing a single result
2. target descriptors from different reference frames are averaged together to form a single set of descriptors, producing a single result
3. registration to each set of target descriptors is performed separately (in parallel), producing distinct results for each reference frame

Multiple results for the same keypoint could be reduced to a single final result by averaging them, selecting one based on lowest final loss, or selecting one based on highest consensus.

While one or more of these methods could produce better results in specific cases, the improvements were not generalizable across different datasets and different combinations of reference frames. Given their significant computational cost, we elected not to include these implementations in ZephIR.

#### Motion prediction

In order to predict the global motion of keypoints between the parent and child frames, ZephIR uses trilinear interpolation with Gaussian blurring to generate a flow field between the two frames. The displacement vectors at the parent keypoint coordinates are sampled from this flow field and used to calculate better initial child keypoint coordinates. In Fig. 9, we evaluate the improvements in the final tracking results as we add more partial annotations to generate more accurate flow fields.

**Figure 9.**
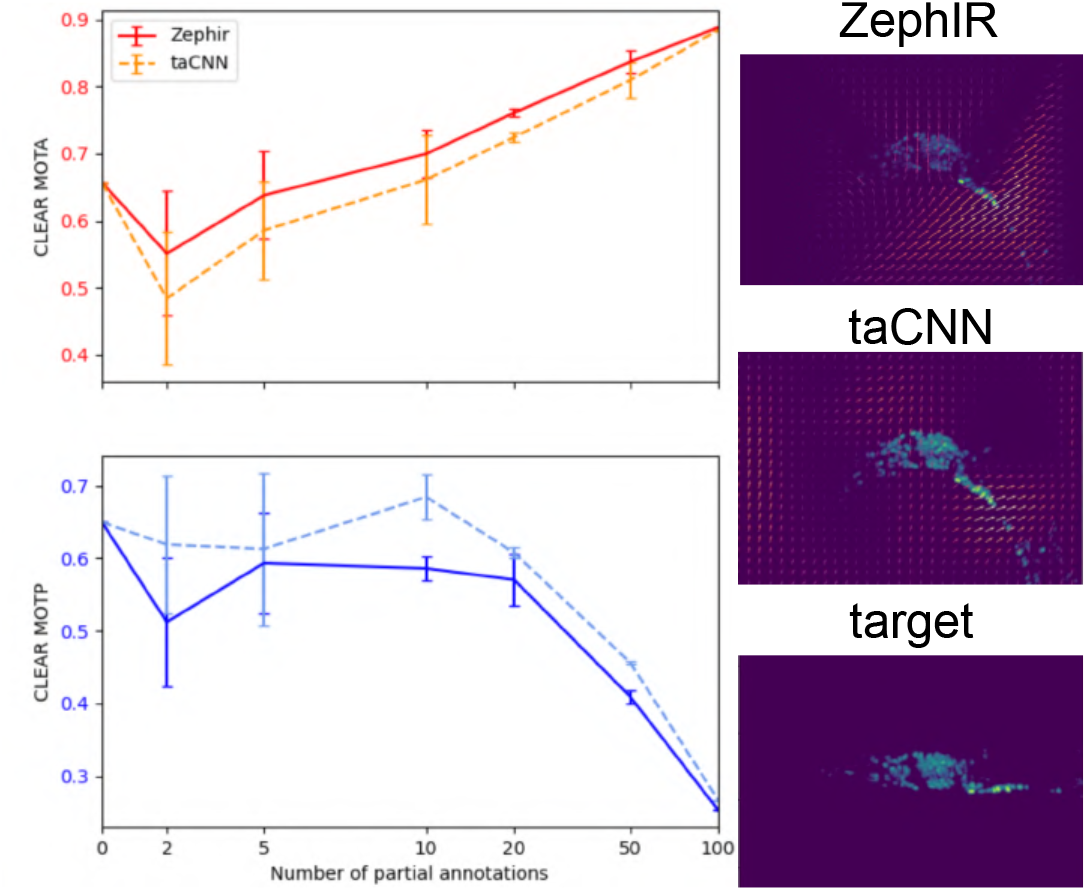
Testing motion prediction. We evaluate tracking accuracy (top left, higher is better) and precision (bottom left, lower is better) for keypoints in a child frame (bottom right) as we add partial annotations. In these tests, we use another frame (top, middle right) as both the parent (providing initial positions for keypoints) and reference frame (providing reference descriptors for registration). We compare the improvements in performance from using displace vectors sampled from ZephIR’s interpolated flow field (top right) and those sampled from taCNN’s low-frequency deformation field (top middle).

In comparison, we test the flow field generation method used in taCNN (26). In this algorithm, neurons are tracked by analyzing images through two different convolutional neural networks. The first CNN produces coarse initial predictions for keypoints in the target image. Displacements between these predictions and a manually annotated frame are used to fit a deformation field while restricting its Fourier modes to low frequencies and regularizing its divergence. The deformation field is used to warp the annotated frame to match the global posture of the target image and generate new training frames to intelligently augment the amount of data available for training the second, more accurate CNN.

We use the same deformation field in place of our flow field to calculate initial keypoint coordinates for the child frame. However, instead of using a CNN to produce the coarse predictions, we use the displacements between the parent frame and the partial annotations in the child frame to fit the deformation field. In Fig. 9, we compare the improvements in accuracy from taCNN’s low-frequency deformation field (dotted line) to that of ZephIR’s method of motion prediction (solid line).

### Additional examples

#### Neurons in deforming

Hydra. Unlike the freely moving *C. elegans, Hydra* does not exhibit clear spatial or postural patterns over time. Few algorithms have been proposed to target such systems. In particular, the EMC2 algorithm (46) has been developed to detect neuron tracklets and use elastic deformation models to link tracklets in freely behaving *Hydra*. However, EMC2 often requires a large number of neurons to build a good elastic deformation model of the posture dynamics, and its accuracy drops for longer videos. In comparison, Fig. 10 illustrates the quality of ZephIR’s tracking of a subset of neurons in behaving *Hydra*, achieving a higher accuracy (88.0%) for this particular dataset over 1000 frames than EMC2 (83.8%).

**Figure 10.**
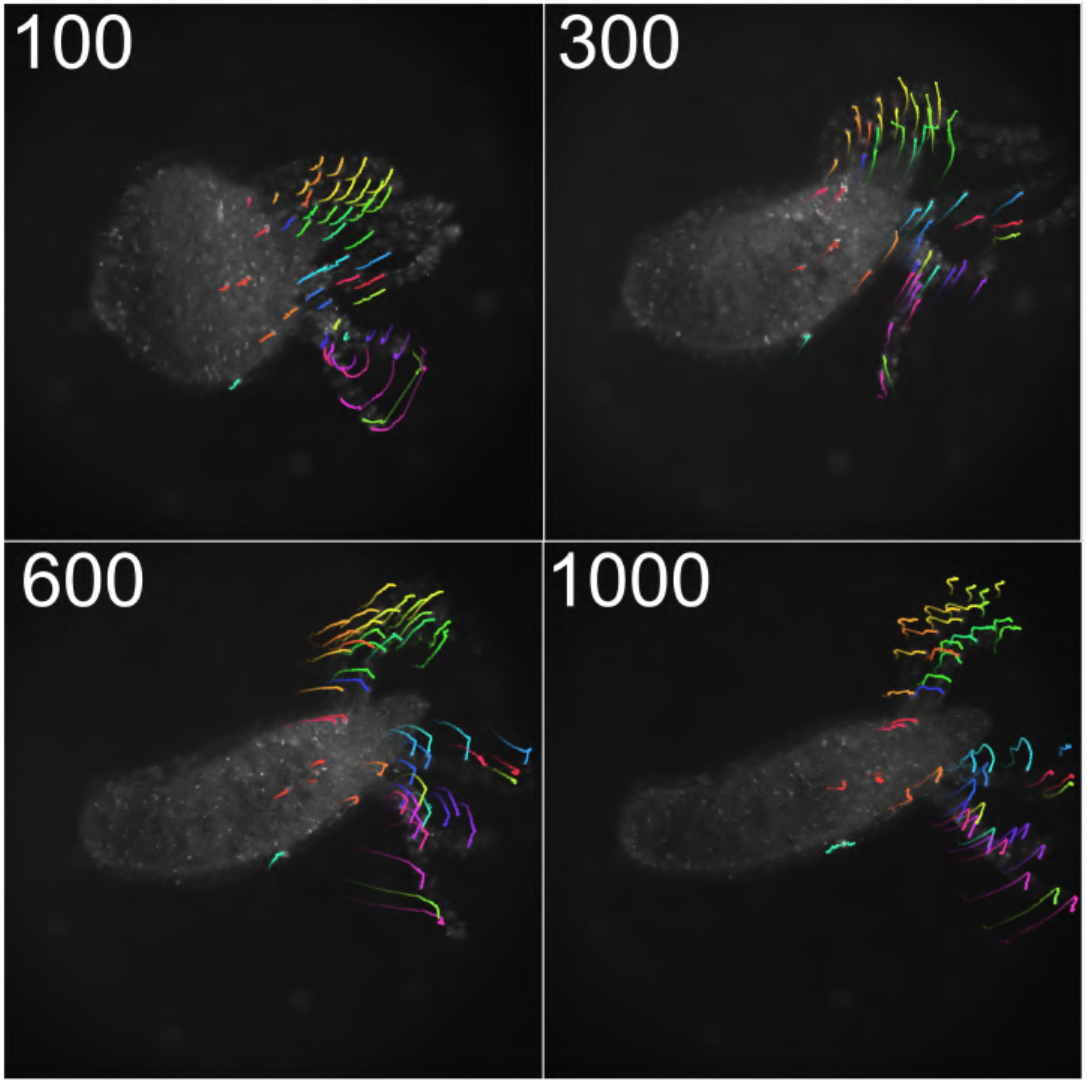
Freely deforming Hydra. We track 50 neurons across 1000 frames with four reference frames. Each panel shows a trail of the neurons’ motions for the 100 frames preceding the frame shown.

### List of user-tunable parameters

- `dataset`: Path to data directory to analyze.
- `load_checkpoint`: Load existing checkpoint.pt file and resume from last run. [*default: False*]
- `load_args`: Load parameters from existing args.json file. [*default: False*]*
- `allow_rotation`: Enable optimizable parameter for rotating image descriptors. This may be helpful for datasets that have clear rotational changes in shape, but is generally superfluous for nucleus tracking. [*default: False*]
- `channel`: Choose which data channel to analyze. Leave out to use all available channels.
- `clip_grad`: Maximum value for gradients for gradient descent. Use −1 to uncap. [*default:* 1.0] *TIP:* If the motion is small, set lower to *≈* 0.1. This is a more aggressive tactic than lr_ceiling.
- `cuda`: Toggle to allow GPU usage if a CUDA-compatible GPU is available for use. [*default: True*]
- `dimmer_ratio`: Coefficient for dimming non-foveated regions at the edges of descriptors. [*default:* 0.1]
- `exclude_self`: Exclude annotations with provenance “ZEIR” which is the current provenance for ZephIR itself. Effectively, this allows you to do the following: if True, you can track iteratively with separate calls of ZephIR without previous results affecting the next; if False, you can use the previous results as partial annotations for the next. [*default: True*]
- `exclusive_prov`: Only include annotations with this provenance.
- `fovea_sigma`: Width of Gaussian mask over foveated regions of the descriptors. Decreasing this can help prioritize keeping a neuron at the center. Increase to a large number or set at −1 to disable. [*default:* 2.5]
- `gamma`: Coefficient for gamma correction. Integrated into
- `get_data`. [*default:* 2]
- `grid_shape`: Size of the image descriptors in the xy-plane in pixels. Increasing this may provide a better view of the neighboring features and avoid instabilities due to empty (only 0’s) descriptors, but it will also slow down performance. [*default:* 25]
- `include_all`: Include all existing annotations to save file, even those ignored for tracking. In the case that annotations and ZephIR results have matching worldlines in the same frame, annotations will override the results. [*default: True*] *TIP:* If using `save_mode`=‘o’, set this argument to “True” to avoid losing any previous annotations. On the other hand, using “False” with `save_mode`=‘w’ may allow you to compare the annotations in “annotations.h5” to the newly saved results in “coordinates.h5”.
- `lambda_d`: Coefficient for feature detection loss, *λ*_*D*_. This regularization is turned on at the last *n_epoch_d* of each optimization loop with everything else turned off. Set to −1 to disable. [*default: ≠*1.0]
- `lambda_n`: Coefficient for spring constant for intra-keypoint spatial regularization, *λ*_*N*_. Spring constants are calculated by multiplying the covariance of connected pairs by this number and passing the result through a ReLU layer. The resulting loss is also rescaled to this value, i.e. it cannot exceed this value. If a covariance value is unavailable, the spring constant is set equal to this number. [*default:* 1.0] *TIP:* Increase up to 10.0 for non-deforming datasets. Decrease down to 0.01 or turn off for large deformation. Optimal value tends to be between 1.0 − 4.0. Set to 0 or −1 if regularization is unnecessary (this can speed up performance).
- `lambda_n_mode`: Method to use for calculating *λ*_*N*_.
  - `disp`: use inter-keypoint displacements
  - `norm`: use inter-keypoint distances (rotation is not penalized)
  - `ljp`: use a Lenard-Jones potential on inter-keypoint distances (collapsing onto the same position is highly penalized.
  - *default: disp*
- `lambda_t`: Coefficient for temporal smoothing loss, *λ*_*T*_, enforcing a 0th-order linear fit for intensity over `n_frame` frames. [*default: ≠*1.0] *TIP:* 0.1 generally matches order of magnitude of registration loss. Increase up to 1.0 for non-deforming datasets. Set to 0 or −1 if regularization is unnecessary (this can dramatically speed up performance). Alternatively, setting `n_frame` to 1 will also disable this.
- `load_nn`: Load in spring connections as defined in *nn_idx*.*txt* if available, save a new one if not. This file can be edited to manually define the connections by worldline ID. The first column is connected to all proceeding columns. [*default: True*] *WARNING:* Note that all connections are necessarily symmetric (i.e. if object0 connected to object2, then object2 must also be connected to object0) even if not defined as such in the file due to how gradients are calculated and accumulated during optimization.
- `lr_ceiling`: Maximum value for initial learning rate. Note that, by default, learning rate decays by a factor of 0.5 every 10 epochs. [*default:* 0.2] *TIP:* If motion is small, set lower to *≈* 0.1. Can use with `clip_grad`, but may be redundant.
- `lr_coef`: Coefficient for initial learning rate, multiplied by the distance between current frame and its parent. [*default:*2.0]
- `lr_floor`: Minimum value for initial learning rate. [*default:* 0.02]
- `motion_predict`: Enable parent-child flow field to predict low-frequency motion and initialize new keypoints positions for current frame. Requires partial annotations for that frame. [*default: False*] *TIP:* Identify and annotate a critical subset of keypoints with large errors. These along with `motion_predict` can dramatically improve tracking quality. Note that this flow field does *not* affect descriptors to avoid distortion or image artifacts.
- `n_chunks`: Number of steps to divide the forward pass into. This trades some computation time to reduce maximum memory required. [*default:* 10]
- `n_epoch`: Number of iterations for image registration, *λ*_*R*_. [*default:* 40]
- `n_epoch_d`: Number of iterations for feature detection regularization, *λ*_*D*_. [*default:* 10]
- `n_frame`: Number of frames to analyze together for temporal loss (see `lambda_t`). Set to 1 if regularization is unnecessary. [*default:* 1]
- `n_ref`: Manually set the number of keypoints. Leave out to set the number as the maximum number of keypoints available in an annotated frame. *WARNING:* This requires at least one annotated frame with exactly `n_ref` keypoints. The ID’s from the first frame with exactly `n_ref` keypoints are used to pull and sort annotations from other annotated frames.
- `nn_max`: Maximum number of neighboring keypoints to be connected by springs for calculating *λ*_*N*_. [*default:* 5]
- `save_mode`: Mode for saving results.
  - `o`: overwrite existing ’annotations.h5’ file *WARNING:* While provenance can ensure manual annotations remain intact and separable from ZephIR results, this can still be volatile! Backup of the existing *annotations*.*h5* is created before saving. Consider enabling `include_all`.
  - `w`: write to a new ’coordinates.h5’ file and replace any existing file
  - `a`: append to existing ’coordinates.h5’ file
  - *default: o*
- `sort_mode`: Method for sorting frames and determining parent-child branches.
  - `similarity`: minimizes distance between parent and child
  - `linear`: branches out from reference frames linearly forwards and backwards, with every parent-child one frame apart, until it reaches the first frame, last frame, or another branch (simplest and fastest)
  - `depth`: uses shortest-path grid search, then sorts frames based on depth in the resulting parent-child tree (this can scale up to *O*(*n*^4^) in computation with number of frames)
  - *default: similarity*
- `t_ignore`: Ignore these frames during registration. Leave out to analyze all frames.
- `t_ref`: Only search these frames for available annotations. Leave out if you want to process all annotations.
- `wlid_ref`: Identify specific keypoints to track by world-line ID (note “worldline ID” and “track ID” are used synonymously). Pulls all available annotations for these keypoints. Leave out to track all available keypoints.
- *WARNING:* This will supercede `n_ref`.
- `z_compensator`: Multiply gradients in the z-axis by (1+ *z*_*compensator*). Since the internal coordinate system is rescaled from -1 to 1 in all directions, gradients in the z-axis may be too small when there is a large disparity between the xy- and z-shapes of the dataset, and thus fail to track motion in the z-axis. Increasing this will compensate for the disparity. Note that gradients will still be clipped to (*clip*_*grad * z*_*compensator*) if `clip_grad` is enabled. Set to 0 or -1 to disable. [*default: ≠*1]

### List of examples with parameter choices and explanations

We tested a number of datasets across various systems. For each dataset, we show the original movie, a movie annotated with the tracking result, a list of parameters used, and a brief explanation for each parameter. Note that only parameters that were changed from the default are listed here.

**Figure 11.**
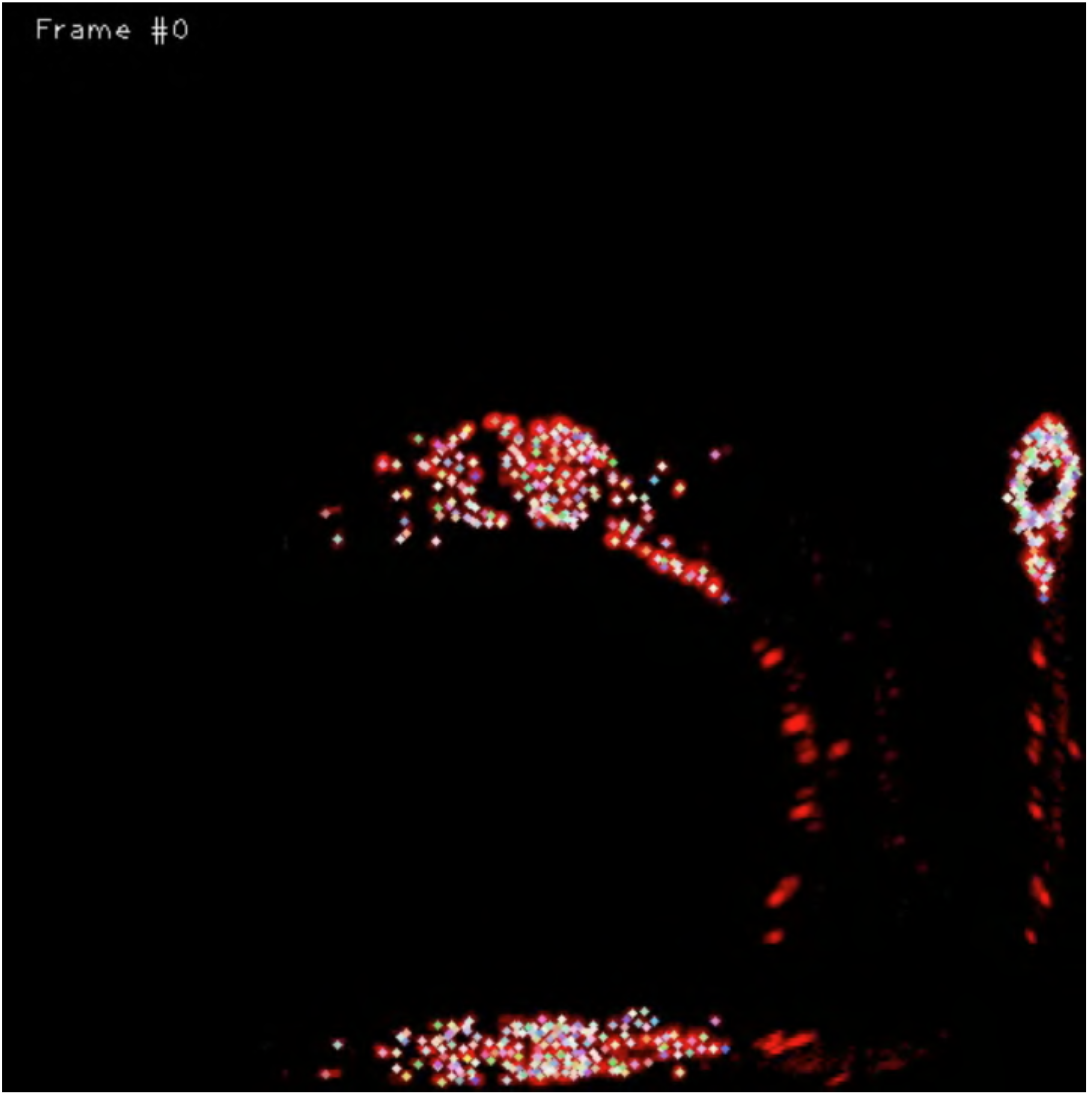
Freely moving *C. elegans*. Movie available at: https://github.com/venkatachalamlab/NeuronIR/blob/main/docs/examples.md

#### Neurons in freely behaving C. elegans

- `channel` = 1: This dataset has 2 channels, but only the second channel has the neurons that we want to track.
- `clip_grad` = −1: This dataset has significant motion in all directions. To accomodate neurons with large displacements between parent and child frames, we disable gradient clipping, relying on learning rates to adjust how much displacement we allow.
- `fovea_sigma` = 10: Along with `grid_shape`, this opens up the view range for descriptors and de-emphasizes the importance of the center of the descriptor relative to its neighbors. While this can be detrimental for rapidly deforming densely-packed clusters, it is particularly helpful for neurons towards the edges of the volume.
- `grid_shape` = 49: This increases the size of the descriptors. Generally, it should be about *≈* 150% of the cell’s size in pixels, but we use much larger descriptors here to avoid having any descriptors with all zeros, which can cause instabilities during gradient descent. This is usually not an issue, but the head swings generate large motions in particularly sparse areas of the volume.
- `lambda_n_mode` = *norm*: This dataset sees significant rotations in relative positions of neighboring neurons. *Norm* mode avoids penalizing those neighbors.
- `lr_ceiling` = 0.1: This limits frame-to-frame displacement of each neuron. Since we uncapped the gradient values, we can be a little more aggressive with this parameter.
- `lr_floor` = 0.01: We lower this to ensure that parent-child frames that are very close together also produce similar neuron positions. We have good, distinct clusters of similar frames around each reference frame, so we can lower this further.
- `motion_predict` = *True*: We verify and fix 10 neurons to use as partial annotations across all frames (see Figure 3).
- To make full use of these partial annotations, we turn on `motion_predict` to improve tracking for the rest of the neurons.
- `z_compensator` = 4.0: This dataset has noticeable motion in the z-axis even when the center of the volume is fixed, but its size in z is 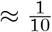 of the xy-shape. We increase this parameter to compensate for the disparity.

**Figure 12.**
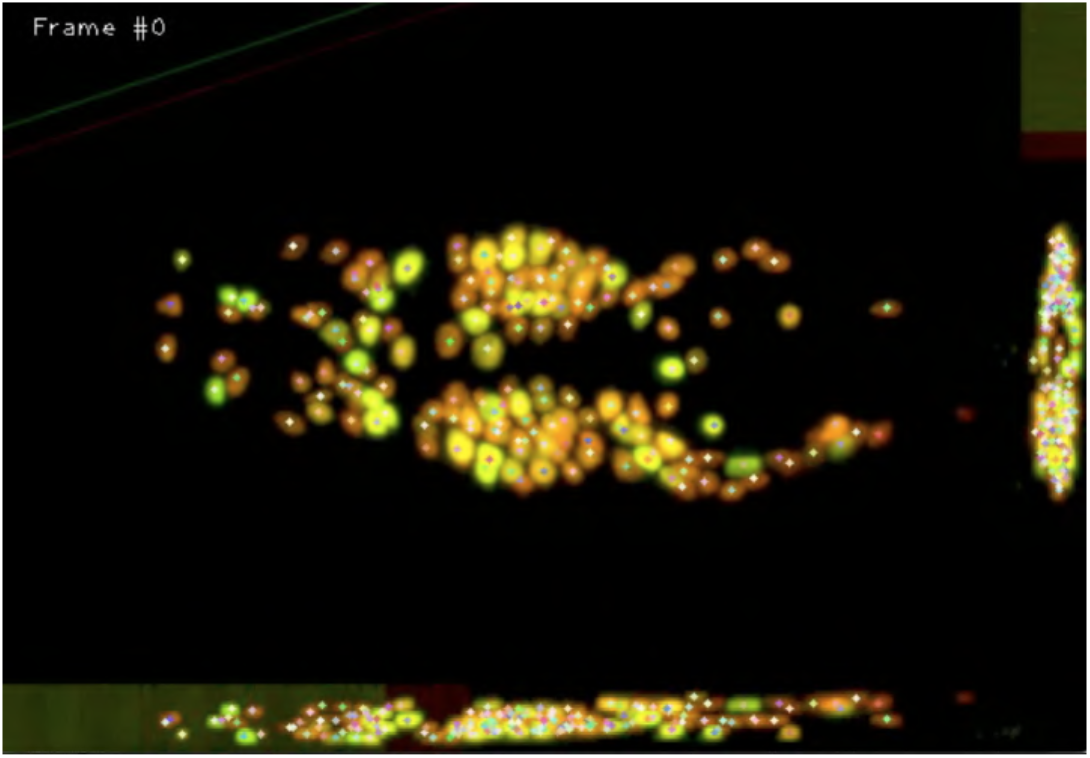
Immobilized unc-13 *C. elegans*. Movie available at: https://github.com/venkatachalamlab/NeuronIR/blob/main/docs/examples.md

#### Neurons in immobilized unc-13 C. elegans

- `clip_grad` = 0.2: Despite the large jumps for some neurons during pumping events, the dataset as a whole does not exhibit much motion. Clipping the gradients prevents tracking results from becoming wildly inaccurate for the low-motion neurons.
- `lambda_n` = 0.2: Most of the neurons here do not show significant motion, but those that do are often isolated and move independently. Lowering this parameter prevents more stationary neurons from being pulled out of position due to nearby high-motion neurons.
- `lr_ceiling` = 0.1: Along with `clip_grad`, this prevents tracking results from moving too much frame-to-frame.
- `lr_floor` = 0.01: Some parent-child pairs do not see any motion at all. We lower this parameter to ensure the neuron positions also do not move for those frames.

#### Chinese hamster ovarian nuclei

- `clip_grad` = 0.33: This dataset exhibits large fluctuations in parent-child frame similarities. We increase the learning rates parameters to accomodate the larger range, but we reduce the gradient values here to prevent tracking results from moving too much.
- `fovea_sigma` = 10: We track very large cells for this dataset. Along with `grid_shape`, this parameter is increased to capture the entire cell in the descriptor.
- `grid_shape` = 125: We track very large cells for this dataset. Along with `fovea_sigma`, this parameter is increased to capture the entire cell in the descriptor.
- `lambda_n` = −1: Cells in this dataset undergo mitosis. ZephIR is built to track a fixed number of keypoints, but we can accomodate the mitosis events by starting from the last frame with the maximum number of cells and allowing keypoints to collapse together as we move backwards. We disable spring connections to avoid penalizing collapsing keypoints.
- `lr_ceiling` = 0.4: This dataset exhibits large fluctuations in parent-child frame similarities. Along with `lr_floor`, we increase this to accomodate the larger range.
- `lr_floor` = 0.06: This dataset exhibits large fluctuations in parent-child frame similarities. Along with `lr_ceiling`, we increase this to accomodate the larger range.
- `sort_mode` = *linear*: Cells in this dataset undergo mitosis. ZephIR is built to track a fixed number of keypoints, but we can accomodate the mitosis events by starting from the last frame with the maximum number of cells and allowing keypoints to collapse together as we move backwards. This means temporal ordering becomes important, so we set this parameter to linear.

**Figure 13.**
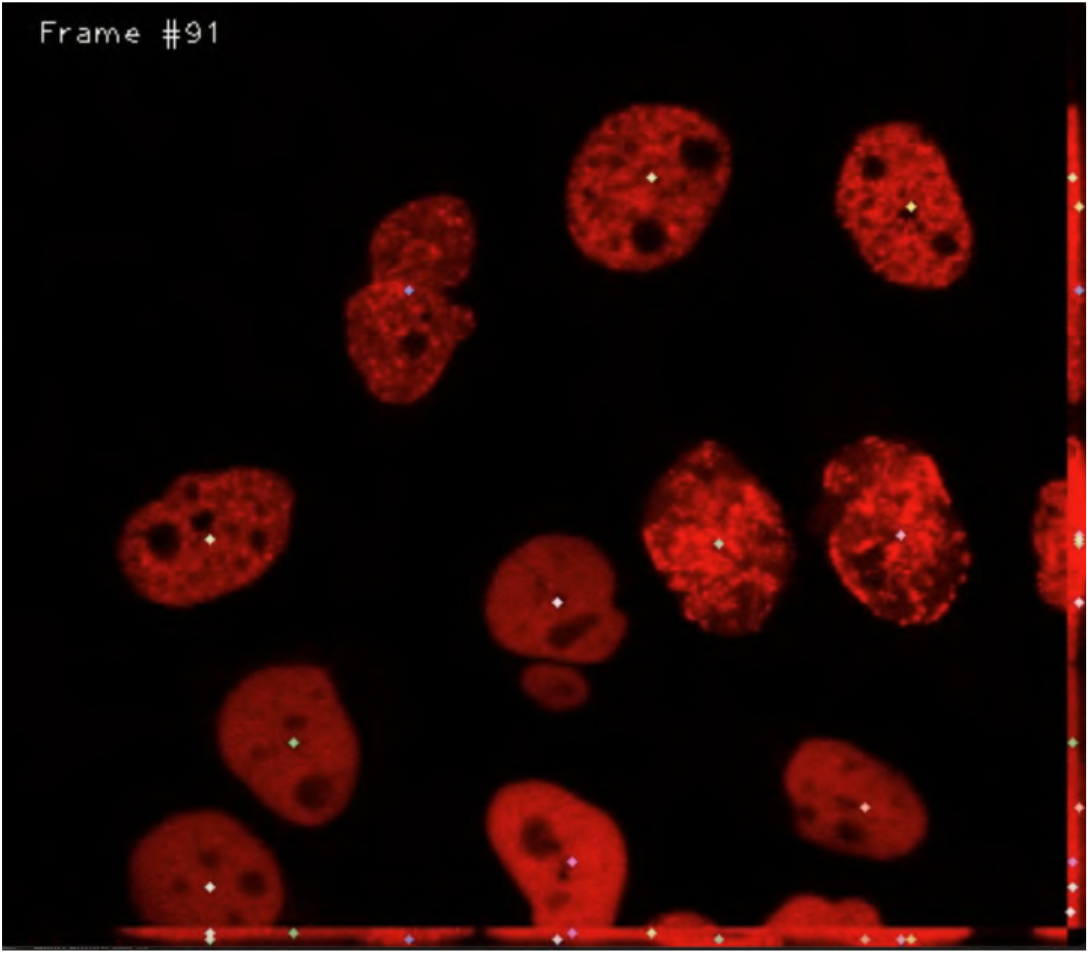
Chinese hamster ovarian nuclei overexpressing GFP-PCNA. Movie available at: https://github.com/venkatachalamlab/NeuronIR/blob/main/docs/examples.md Data available at: http://celltrackingchallenge.net/3d-datasets/

#### Neurons in Hydra

- `allow_rotation` = *True*: Features in this dataset show clear rotation, especially in the tentacles. To accomodate this, we enable optimization for an additional parameter that controls rotation of descriptors in the xy-plane.
- `grid_shape` = 11: Each neuron in this dataset is small, so a lower value here is sufficient.
- `lambda_n` = 0.1: While the tentacles create a good biological scaffolding for the neurons, they deform and stretch over time. We lower the spring constants to accomodate these deformations.
- `lambda_n_mode` = *norm*: This dataset shows clear rotations in relative positions of neighboring neurons, especially in the tentacles. *norm* mode avoids penalizing those neighbors.
- `lr_floor` = 0.08: The large, bright body obscures the changes in the tentacles when calculating parent-child frame similarities. We increase this parameter to compensate.
- `sort_mode` = *linear*: This dataset does not repeatedly sample the same postures, but rather continuously deforms over time. To reflect this, we track linearly.

**Figure 14.**
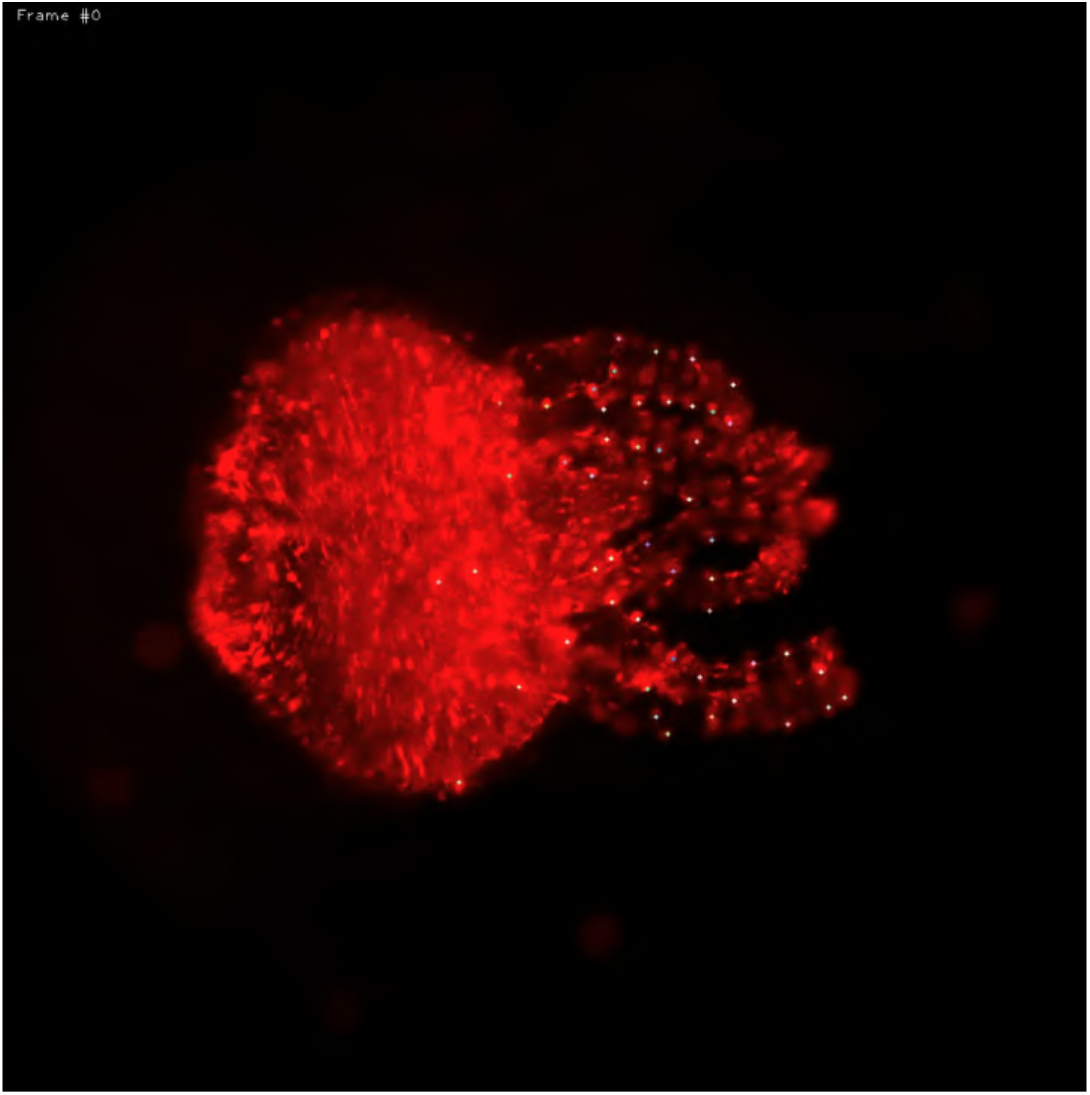
Freely moving *Hydra*. Movie available at: https://github.com/venkatachalamlab/NeuronIR/blob/main/docs/examples.md Data available at: https://www.ebi.ac.uk/biostudies/studies/S-BSST428

#### Body parts of a behaving mouse

- `allow_rotation` = *True*: Features in this dataset show clear rotation, especially in the paws. To accomodate this, we enable optimization for an additional parameter that controls rotation of descriptors in the xy-plane.
- `dimmer_ratio` = 0.8: Unlike fluorescent microscopy data, this is a particularly feature-rich dataset. Increasing this parameter emphasizes the neighboring features relative to the centers of descriptors.
- `fovea_sigma` = 49: We track very large features for this dataset. Along with `grid_shape`, this parameter is increased to capture the entire body part in the descriptor.
- `grid_shape` = 65: We track very large features for this dataset. Along with `fovea_sigma`, this parameter is increased to capture the entire body part in the descriptor.
- `lambda_n` = 0.1: While the skeleton creates a good biological scaffolding for the mouse, this dataset lacks a third dimension and thus the distances between body parts are not well-preserved in the image. We lower the spring constants to accomodate this.
- `lambda_n_mode` = *norm*: This dataset shows clear rotations in relative positions of neighboring features, especially in the paws. *norm* mode avoids penalizing those neighbors.
- `nn_max` = 3: We only track 10 keypoints for this dataset. We reduce the number of maximum neighbors to reflect the small number of total points.

**Figure 15.**
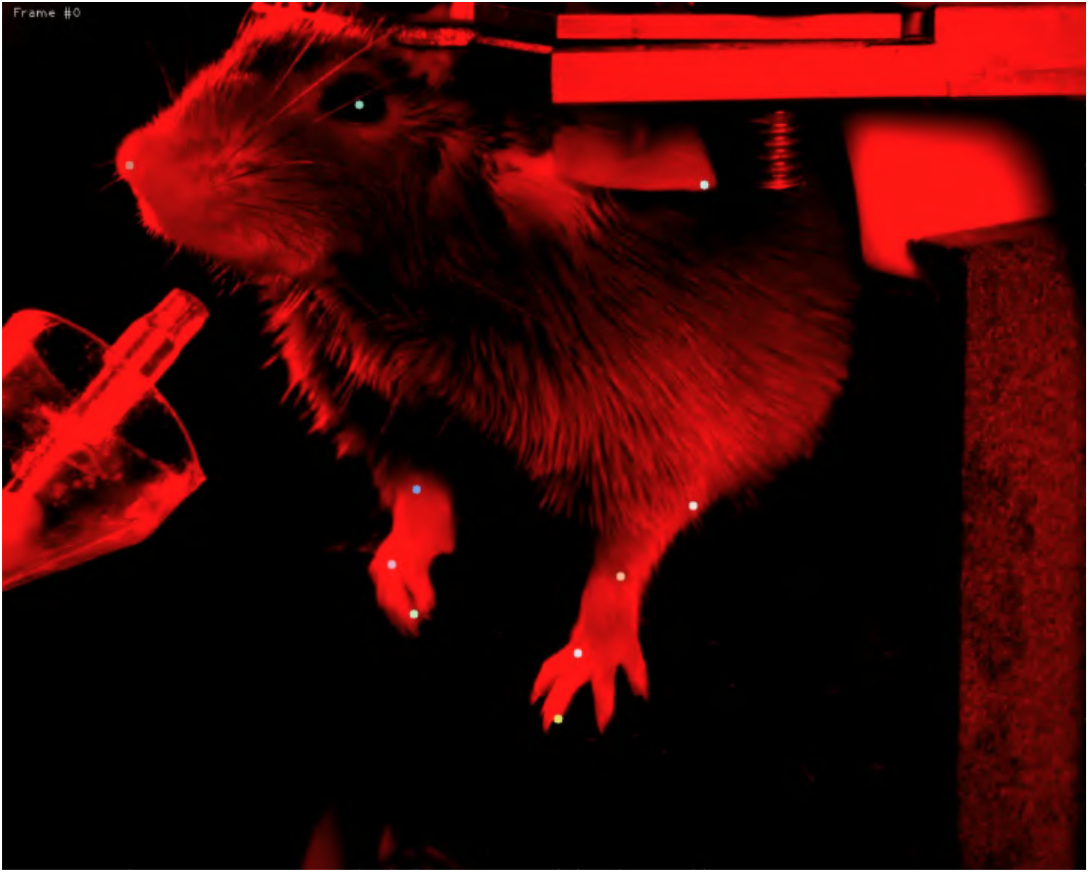
Behaving mouse. Movie available at: https://github.com/venkatachalamlab/NeuronIR/blob/main/docs/examples.md Data available at: https://ibl.flatironinstitute.org/public/churchlandlab/Subjects/CSHL047/2020-01-20/001/raw_video_data/

